# Pdgfrβ signaling orchestrates meningeal repair via the mobilization of arachnoid cells

**DOI:** 10.64898/2026.05.13.723714

**Authors:** Payel Banerjee, Michael Demarque, Axel Benchetrit, Dorian Champelovier, Cynthia Froc, Manon Paul, Hannah Wiggett, Arnim Jenett, Jean-Pierre Levraud

## Abstract

Zebrafish possess remarkable regeneration abilities, including the capacity to repair their central nervous system (CNS). Leveraging the optical accessibility of the zebrafish brain, we investigated the mechanisms underlying meningeal repair following laser-induced brain injuries to the optic tectum. In previous work, using live imaging of laser-induced surface injuries to the optic tectum in juvenile zebrafish, we identified a population of flat PDGFRβ^+^ cells that rapidly migrate to the wound site and contribute to meningeal repair. Here, using pharmacological inhibition or genetic perturbation of PDGFRβ, we show that recruitment of these PDGFRβ^+^ meningeal cells is strongly dependent on PDGFRβ signaling, unlike recruitment of PDGFRβ^+^ pericytes deeper in the wound. Furthermore, PDGFRβ inhibition diminishes neurite regrowth and macrophage recruitment. Using photoconversion assays, we traced the origin of PDGFRβ^+^ meningeal cells that migrated to the wound in response to injury, in the midbrain-forebrain and midbrain-hindbrain sulci. Our findings highlight PDGFRβ’s pivotal role in orchestrating meningeal repair and reveal novel cellular dynamics during CNS regeneration. These results provide insights into potential therapeutic strategies for enhancing brain repair and mitigating fibrosis in mammals, where meningeal scarring remains a barrier to CNS regeneration.

**Highlights:**

PDGFRβ signaling orchestrates the rapid accumulation of meningeal cells at CNS injury sites to drive tissue remodeling.
PDGFR activity acts as a critical regulator, co-recruiting PDGFRβ^+^ meningeal cells and mfap4+ macrophages to the lesion.
Meningeal cells are essential for neural repair, with PDGFR inhibition leading to significantly reduced neurite density.
Zebrafish maintain locomotor resilience post-injury, identifying a clear distinction between meningeal-driven axonal regrowth and basic motor circuitry recovery.
Establishes a high-resolution zebrafish platform to identify conserved meningeal targets for mammalian CNS regeneration.

## Introduction

The regenerative capacity of the central nervous system (CNS) represents a fundamental divergence in vertebrate evolution. In mammals, traumatic CNS injury typically results in permanent functional deficits, driven by a complex inflammatory cascade and the formation of a growth-inhibitory fibrotic scar^1^. This scarring process is largely orchestrated by heterogeneous fibroblast populations and pericyte-derived myofibroblasts that deposit a dense, inhibitory extracellular matrix (ECM)^2^. Conversely, zebrafish (Danio rerio) exhibit an extraordinary ability for scar-free CNS repair, efficiently restoring both tissue architecture and neural function. Understanding the mechanisms that allow zebrafish to maintain a “permissive” environment rather than a fibrotic one, is essential for developing regenerative strategies for mammalian CNS trauma.

Successful CNS regeneration is increasingly understood as a multi-lineage “neurovascular orchestration”, where non-neuronal cells actively guide the timing and spatial organization of tissue repair^3,4^. A primary molecular driver of this process is the platelet-derived growth factor receptor beta (Pdgfrβ) signaling pathway. While PDGFRβ is a well-established regulator of mural cell recruitment and vascular integrity^5^, recent studies have expanded its role to broader regenerative functions. In the zebrafish spinal cord, a critical switch in PDGFRβ^+^ cell-derived ECM composition prevents inhibitory scarring and promotes axonal bridging^2^. Similarly, Pdgfr signaling is required for re-vascularization and the modulation of the immune landscape in the regenerating retina^6^ and for epicardial function during heart regeneration^7^. However, the specific mechanistic contribution of PDGFRβ+ cells within the meninges, the protective barrier of the brain, remains poorly defined.

The meninges are now recognized as a dynamic hub for CNS homeostasis^8^, harboring specialized populations that interact to regulate angiogenesis, waste clearance, and immune surveillance^9,10,11^. Within this niche, mural lymphatic endothelial cells (muLECs) and their supporting cells govern the formation and maintenance of the intracranial environment^12,13^. Following injury, prompt meningeal reconstruction is a mechanical and physiological prerequisite for neural recovery, involving scaffolding proteins like AKAP12 to mediate initial repair^14^.

We previously established a juvenile zebrafish brain injury model using biphoton-generated UV irradiation to target the superficial layers of the optic tectum^15^. By examining 21 days post-fertilization (dpf) larvae, a stage where the meninges have attained structural complexity, we identified a unique population of flat, PDGFRβ^+^ arachnoid-like cells. These cells reside in specialized “reservoirs” within the cranial sulci and migrate toward the injury site to re-establish the meningeal barrier^16,17,18^.

In the present study, we demonstrate that the recruitment and migration of these PDGFRβ^+^ arachnoid cells are strictly governed by PDGFRβ signaling. Utilizing pharmacological inhibition and Kaede-based photoconversion assays, we show that disrupting PDGFRβ activity not only halts meningeal repair but also significantly impairs the wider regenerative niche. This impairment is characterized by a secondary failure in macrophage accumulation and a marked reduction in neurite density at the lesion site. These findings suggest that PDGFRβ^+^ meningeal cells act as essential coordinators of the repair process, influencing both immune response and neural regrowth. By defining the molecular drivers of meningeal reconstruction, this work provides a framework for understanding why mammalian CNS repair fails and identifies Pdgfrβ as a potential therapeutic target for modulating the fibrotic response to injury^15,19^.

## Materials and Methods

### Zebrafish Husbandry and Transgenic Lines

Eggs from natural spawning were bleached and embryos incubated at 28°C in embryonic medium (EM) following established protocols^20^. Wild-type (WT) and casper zebrafish lines were obtained from the Zebrafish International Resource Center (ZIRC, Eugene, OR, USA) or the European Zebrafish Resource Center (EZRC, Karlsruhe, Germany). Environmental conditions were maintained as follows: a photoperiod of 14 hours light and 10 hours dark, temperature of 26.5 ± 1°C, pH of 7.8 ± 0.1, conductivity of 240 ± 30 µS/cm, ammonia (NH₄⁺) at 0 mg/L, nitrite (NO₂⁻) at 0 mg/L, and nitrate (NO₃⁻) below 50 mg/L. Fish were initially fed rotifers (Brachionus plicatilis, approximately 500 per fish per day) for the first two weeks, followed by a combination of brine shrimp (*Artemia nauplii*, approximately 250 per fish per day) and dry food (Skretting Gemma Micro) twice daily. All protocols were approved by the local ethics committee for animal experimentation (CEEA 59) and the French Ministry of Research and Education. All procedures were performed in accordance with European Union Directive 2011/63/EU, with the approval of the local ethics committee (no. 59 CEEA). To visualize and functionally manipulate specific cell populations, we utilized a suite of transgenic reporter and effector lines. Meningeal cell dynamics were tracked using TgBAC(*pdgfrb*:Gal4FF)^ncv24,5^ in combination with either Tg(5xUAS:eGFP)^nkuasgfp1a,21^ or the photoconvertible Tg(5xUAS:Kaede)^rk8,22^. For structural and immune context, we employed Tg(*kdrl*:DsRed)s896^22^ to label the vasculature, Tg(*mfap4*:mCherry-F)^ump6,23^ for macrophages, and Tg(*Xla.Tubb2:*mapple*-*CAAX)^24^ for neurites. To specifically perturb signaling, we used the heat-shock inducible Tg (*hsp70l*:dnpdgfrb-mCherry)^mps2,2^ line to express a dominant-negative version of the Pdgfrβ receptor. All transgenic lines were maintained in either the nacre (*mitfa*^w2/w2^)^25^ or casper (*mitfa*^w2/w2^;*mpv17*^a9/a9^)^26^ mutant backgrounds to ensure optical clarity for live imaging. Experimental animals were generated by crossing the reporter lines described above with *casper* wild-type fish, resulting in progeny with reduced or absent pigmentation (either *nacre* or *casper* phenotype) suitable for deep-tissue confocal microscopy.

### Targeted genetic removal of melanophores through CRISPR-Cas9-mediated editing

To facilitate high-resolution live imaging by eliminating interference from endogenous pigmentation, we generated a transparent nacre phenotype in the experimental transgenic backgrounds. This was achieved by CRISPR/Cas9-mediated targeting of mitfa (microphthalmia-associated transcription factor a), a bHLH-Zip transcription factor essential for the specification and differentiation of neural crest cells into melanophores^27^. Targeting was performed using a multiplexed ribonucleoprotein (RNP) approach. Synthetic crRNAs and tracrRNA (Integrated DNA Technologies) were denatured and hybridized by incubation at 95°C for 5 min, followed by slow cooling to room temperature (25°C) to form stable sgRNAs (sequences provided in Table 1). To form the RNP complexes, sgRNAs were incubated with SpCas9 HiFi protein (Museum National d’Histoire Naturelle) at 37°C for 10 min, reaching a concentration of 13.33 µM sgRNA and 10 µM Cas9 per guide. The individual RNP duplexes were combined and loaded into heat-elongated glass capillaries (dimensions: 1.0 OD x 0.58 ID x 100 L; Harvard Apparatus). One-cell stage embryos—derived from TgBAC(pdgfrb:Gal4FF)^ncv24^;Tg(5xUAS:EGFP)^nkuasgfp1a^ and Tg(hsp70l:dn-pdgfrb)^mps2^ crosses were aligned in 1.5% agarose-embryo medium molds. Each embryo was microinjected with approximately 2 nL of the multiplexed RNP mix, resulting in a final concentration of 6.67 µM per sgRNA and 5 µM per Cas9. Successful biallelic targeting was confirmed by the loss of melanophore pigmentation (the nacre phenotype) at 5 days post-fertilization (dpf) under a binocular magnifying glass. Only larvae exhibiting a complete loss of pigment were selected and reared until 21 dpf for subsequent injury experiments.

**Table 1.**
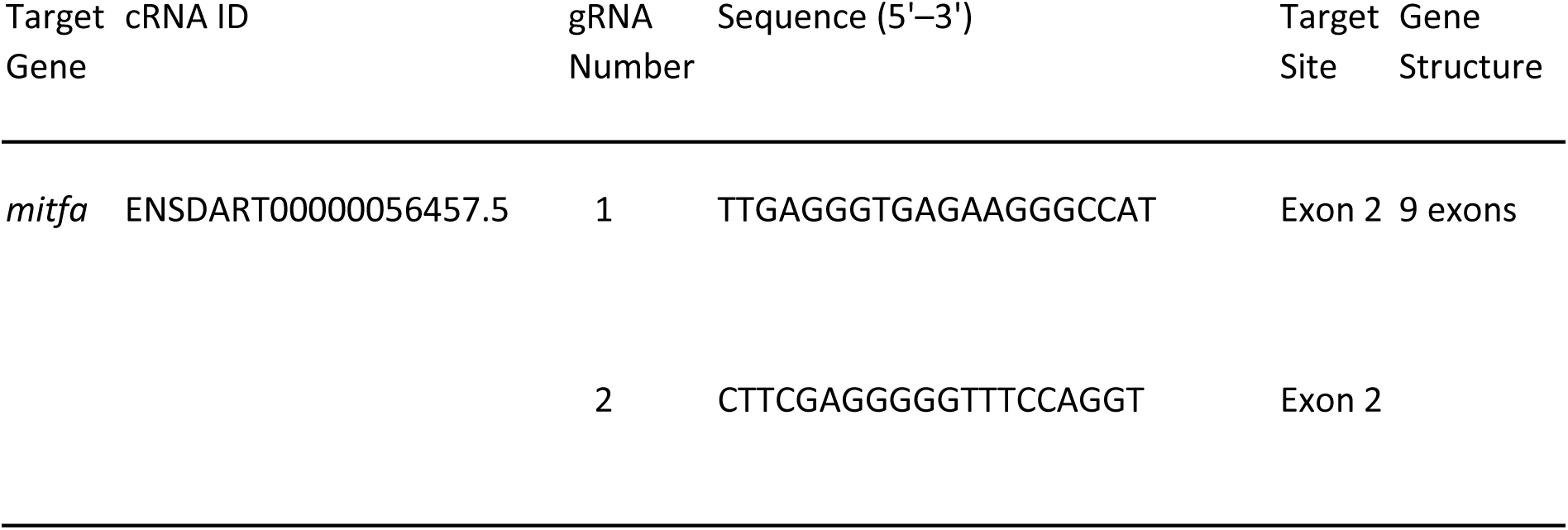
sgRNA sequences for mitfa mutagenesis.

### Drug preparation and treatment

Drug treatments followed the schematic timelines outlined for each experiment. AG1295 (Sigma-Aldrich, Cat# 658550) was dissolved in DMSO (VWR, Catalog: A3672.0100) to prepare a 300 mM stock solution and was used at a final working concentration of 30 µM^28^. As a control, an equivalent volume of DMSO corresponding to the amount used for the 30 µM AG1295 treatment was administered.

Juvenile larvae were treated with AG1295 from 21 to 22 days post-fertilization (dpf). Live imaging was conducted with the drug present in agarose and embryo water supplemented with tricaine. To preserve DMSO’s and AG1295’s light-sensitive properties, the larvae were kept in darkness throughout the treatment and imaging process.

### Preparation of human Platelet-Derived Growth Factor-BB stock solution and PDGF-BB injection

PDGFRβ ligand PDGF-BB treatments were performed as previously described with the following modifications^7,28^. PDGF-BB (Sigma-Aldrich, Cat# P-3201) was dissolved in a 1X PBS buffer to prepare a 1 ng/nl stock solution. As a control, an equivalent amount of BSA was used.

A dose of 180 ng of PDGF-BB or BSA per brain was administered via intracranial microinjection into the midbrain-hindbrain region to ensure delivery of the ligand into the cerebrospinal fluid. Juvenile larvae were treated at 4 days post-injury when they were 25 dpf (days post-fertilization) and imaged live next day, at 26 dpf. Larvae were prepared for injection following the same protocol described in the live imaging section below. After microinjection, fish were released from LMA (low melting agarose), revived in fresh aerated tank water, and then returned to facility tanks until the next imaging session.

### Optic Tectum (OT) Injury

For anesthesia, a 10x stock solution of ethyl 3-aminobenzoate methanesulfonate (MS-222) was prepared by dissolving 400 mg tricaine powder and 800 mg sodium bicarbonate in 200 mL EM^20^. Prior to mounting, 21 days post-fertilization (dpf) zebrafish were anesthetized by immersion in 0.5× MS-222 solution for 45 to 60 seconds until swimming ceased. Zebrafish juveniles, with an average size of 7.8 mm, were mounted in agarose plates containing wells shaped to fit their bodies. These wells were created using 3D-printed molds pressed into 1% agarose. To further immobilize the fish, a drop of 1.5% low-melting point agarose (LMA) was applied at the tail end, which solidified within approximately 10 seconds at room temperature. During the injury procedure, the plate was filled with EM containing 0.5× MS-222, and zebrafish were carefully oriented dorsally using plastic Pasteur pipettes and paintbrushes. After injury, fish were transferred to fresh EM, and air bubbles were introduced with a pump to facilitate recovery. Recovered fish displayed normal swimming and feeding behaviors. Optic tectum surface injuries were induced using a Leica TCS SP8 MP multiphoton microscope equipped with a 25× objective (NA 0.95). Injury was performed in a parallelepiped volume measuring 88 × 88 × 100 µm (x, y, z) below the brain surface. The biphoton laser was tuned to a wavelength of 750 nm with the following settings: power of 2.5 mW, intensity of 50%, and offset of 45%. Injuries were performed with a 2 µm z-step in bidirectional scanning mode and a pixel size of 2 µm. Following injury, fish were released from LMA, revived in fresh aerated tank water, and then returned to facility tanks until the next imaging session^15^.

### Live imaging, image processing and quantification

For live imaging, juvenile zebrafish were anesthetized in embryo medium (EM) containing 0.5× MS-222 and mounted in 1.5% low-melting agarose (LMA) within dental silicone molds. During imaging, they remained submerged in EM with 0.5× MS-222. Imaging was performed using a Leica SP8 confocal microscope equipped with a 25× water immersion objective. Image acquisition parameters included a pixel size of 1.16 × 1.16 µm in the x and y axes, and 2 µm in the z axis, with stacks measuring approximately 300 µm in depth. Each session lasted 25–30 minutes to minimize stress and ensure survival, with imaging typically completed within 10 minutes. Injured areas were identified by absence of any fluorescence protein expression at the center of OT followed by laser injury to ensure consistent imaging over successive time points. After imaging, juveniles were revived and returned to their tanks following the recovery protocol described earlier.

To avoid over- or under-saturation of the sensor, acquisition parameters were calibrated at each time point by saturating the true signal and then reducing it to remove saturation. This calibration was applied to all samples at the given time point, ensuring full emission range capture without compromising subsequent threshold-based analyses.

Image analysis was performed using FIJI software^29^. Fluorescent images were normalized (e.g., flipping, cropping, and denoising) using a custom macro script recorded and adjusted in FIJI. After normalization, a threshold check determined the optimal strategy for each channel, and all images were thresholded uniformly. The final output was the percentage of the positive pixel area normalized to the constant area of the image, focusing on the injured brain surface. The same region in the uninjured hemisphere of the optic tectum (OT) was used as a control.

For each transgenic line used, the percentage area covered by GFP, mApple, DsRed, or mCherry fluorescence was quantified. Statistical analysis, including ANOVA or two-tailed unpaired t-tests, was used to evaluate differences between means.

### Immunohistochemistry

We adapted previously described transperisation and staining protocols for use in 21 dpf juveniles ^15,30,31,32^. The method was refined to balance effective penetration of the overlying dermal and cartilaginous tissues with the preservation of the superficial meningeal cell populations. These adaptations preserved cell integrity and minimized background staining. Immunohistochemistry (IHC) was performed using a higher-than-usual concentration of saponin to facilitate antibody penetration through the skin. Additionally, high antibody dilutions and short incubation times were employed to reduce background staining. The 21 dpf zebrafish were euthanized by immersion in 10× MS-222 (tricaine) dissolved in ice cold embryonic medium. Samples were fixed by incubation in freshly prepared 4% paraformaldehyde (PFA) in PBS for 2 hours at room temperature (RT). Fixation was followed by a brief rinse with 0.1% Tween-20 in PBS (PBST) and six washes (5 minutes each) in PBST with gentle shaking at RT. Samples were blocked for 2 hours at RT in blocking solution containing 10% normal goat serum (NGS), 10% dimethyl sulfoxide (DMSO), 1% Triton X-100, 0.1% Tween-20, and 0.5% saponin in PBS. Primary antibodies were diluted in fast staining solution (2% NGS, 20% DMSO, 0.1% Triton X-100, 0.1% Tween-20, 10 µg/mL heparin, 0.5% saponin in PBS) and incubated with the samples overnight at RT with gentle shaking. Antibodies used included anti-GFP (1:300) and anti-DsRed (1:300). After primary antibody incubation, samples were washed six times in PBST (10 minutes each) at RT with gentle shaking. Secondary antibodies, diluted in fast staining solution, were applied for 24 hours at 4°C with gentle shaking. Subsequently, samples underwent six additional washes in PBST (10 minutes each). To label nuclei, samples were incubated with DAPI (1:1000 dilution in fast staining solution) for 24 hours at RT with gentle shaking. Throughout the process, samples were protected from light to prevent photobleaching. Finally, samples were washed three times in PBST (10 minutes each) at RT with gentle shaking. For imaging, samples were transferred to a high-refractive-index clearing medium (RI = 1.457). The clearing medium was prepared by dissolving 225 g sucrose, 100 g nicotinamide, 50 g triethanolamine, and 500 µL Triton X-100 in double-distilled water to a final volume of 1 L. Samples were progressively immersed in 25%, 50%, 75%, and 100% clearing medium for 30 minutes at each concentration with gentle shaking. Cleared samples were imaged using a Leica laser-scanning confocal microscope equipped with 25× objectives for high-resolution imaging. Image analysis was performed using Amira (2019) and ImageJ software.

### Heat shock treatments

Heat shocks treatment was performed as previously described^4^ and according to the schematic timelines shown with each experiment. For heat shocks, larvae were kept in 50 ml conical centrifuge tubes filled with 40 ml E3 medium^33^, which was placed in a 38°C incubator. Heat shocks of transgenic animals and non-transgenic sibling controls were performed for 1 h at 38°C after which larvae were returned to 28°C.

### Photoconversion and confocal imaging

Photoconversion was carried out on a Leica SP8 confocal microscope, utilizing parameters adapted from previously described methods^22^. Briefly, Tg(*pdgfrb:*Gal4;UAS:Kaede)*; nacre* embryos were maintained in the animal facility at 28 °C until 21 days post-fertilization (dpf). Larvae were then mounted in 1.5% low-melting-point agarose (LMA) for further manipulation. A surface brain injury was induced using a multiphoton system, as described above. Following injury, photoconversion of Kaede protein in a selected region either the telencephalon–optic tectum (OT) border or the midline area, was performed using 405 nm laser excitation and a 25×/0.95 NA water immersion objective. Confocal images were acquired before injury, immediately after injury, and following photoconversion using the same microscope setup.

To detect the green form of Kaede before photoconversion, a 488 nm laser was used. A zoom factor of 5× was applied to highlight the region of interest (ROI), and a Z-stack spanning 50 µm was used to. photoconvert the ROI using 405 nm laser excitation, resulting in the conversion of green Kaede fluorescence to red. Final imaging was carried out in sequential mode using 488 nm and 561 nm lasers to detect green and red Kaede, respectively, at all key time points (pre-injury, post-injury, and post-photoconversion). After imaging, larvae were gently dismounted from the agarose and returned to their tanks in the fish facility upon recovery from anesthesia.

### Spontaneous locomotion assays

We recorded the locomotion of individual larvae from day 0 (one day before performing injury) to 3 days post injury (dpi). The larvae were aged from 20 dpf to 24 dpf. The injury was performed as mentioned previously^15^. Following injury, larvae were released from LMA, revived in fresh aerated tank water, and then returned to fresh 12 well plate. Larvae were placed with 2 ml of EM, supplemented with DMSO or inhibitor, 4 hours before the recordings for habituation. The plates were then placed on an infrared floor, under an infrared-sensitive camera (Zebrabox^34^, Viewpoint) for 30-min recording sessions. Locomotor activity was recorded using ZebraLab software (Videotrack; ViewPoint Life Sciences). At the end of the recording session, the larvae were placed in a freshly renewed medium supplemented with either DMSO or inhibitor in the incubator until the next day’s recording. Each day, plates and medium were renewed before and after each recording session.

### Statistics and reproducibility

Statistical analysis was performed using GraphPad Prism 9. Normality was assessed using the Shapiro-Wilk (or D’Agostino-Pearson) test. For comparisons between two groups, statistical significance was determined using a two-tailed Student’s t-test for normally distributed data and the Mann–Whitney U test for non-normally distributed data. For comparisons among multiple groups, one-way ANOVA was employed for parametric data, while the Kruskal-Wallis test was used for non-parametric data. Data are presented as mean ± SEM. To account for exact p-values below p < 0.0001; For t-tests: t-scores and degrees of freedom (DF) were included in the source data.; For ANOVA: Sums of squares (SS), mean squares (MS), DF, and F-distribution values were provided. For confocal analysis of Tg embryos, expression patterns were confirmed in at least six independent animals.

## Results

### A population of PDGFRβ^+^ cells contribute to meninge repair after surface injury of the zebrafish midbrain

To study the mechanisms of brain and meninges repair, we performed laser-induced injury of the optic tectum surface of 21dpf transgenic zebrafish Tg(*pdgfrb*:gal4; UAS:GFP; *kdrl*:DsRed) in the transparent casper background (Figure 1a,b). As described in our previous work^15^, live imaging of wounded fish revealed, as early as 6 hours post injury (hpi) a population of GFP+ (PDGFRβ^+^) cells present over the wounded area but not on the control side, becoming part of the regenerating meningeal layer (Figure 1c,d; movie S1). This robust meningeal response is readily quantifiable at 24 hpi by measuring GFP+ pixel coverage on a region corresponding to the wounded area (Figure 1e). By contrast, the DsRed+ pixel coverage is reduced at the wounded side (Figure 1f), indicating that laser-induced blood vessel damage is not fully repaired yet. The GFP signal is mostly due to flat GFP+ cells in the meningeal layer above the tectal neuropile; a few GFP+ pericytes inside the parenchyme also contribute to this signal, but not in higher numbers at the wound than on the control side (Figure 1 g,h).

**Figure 1.**
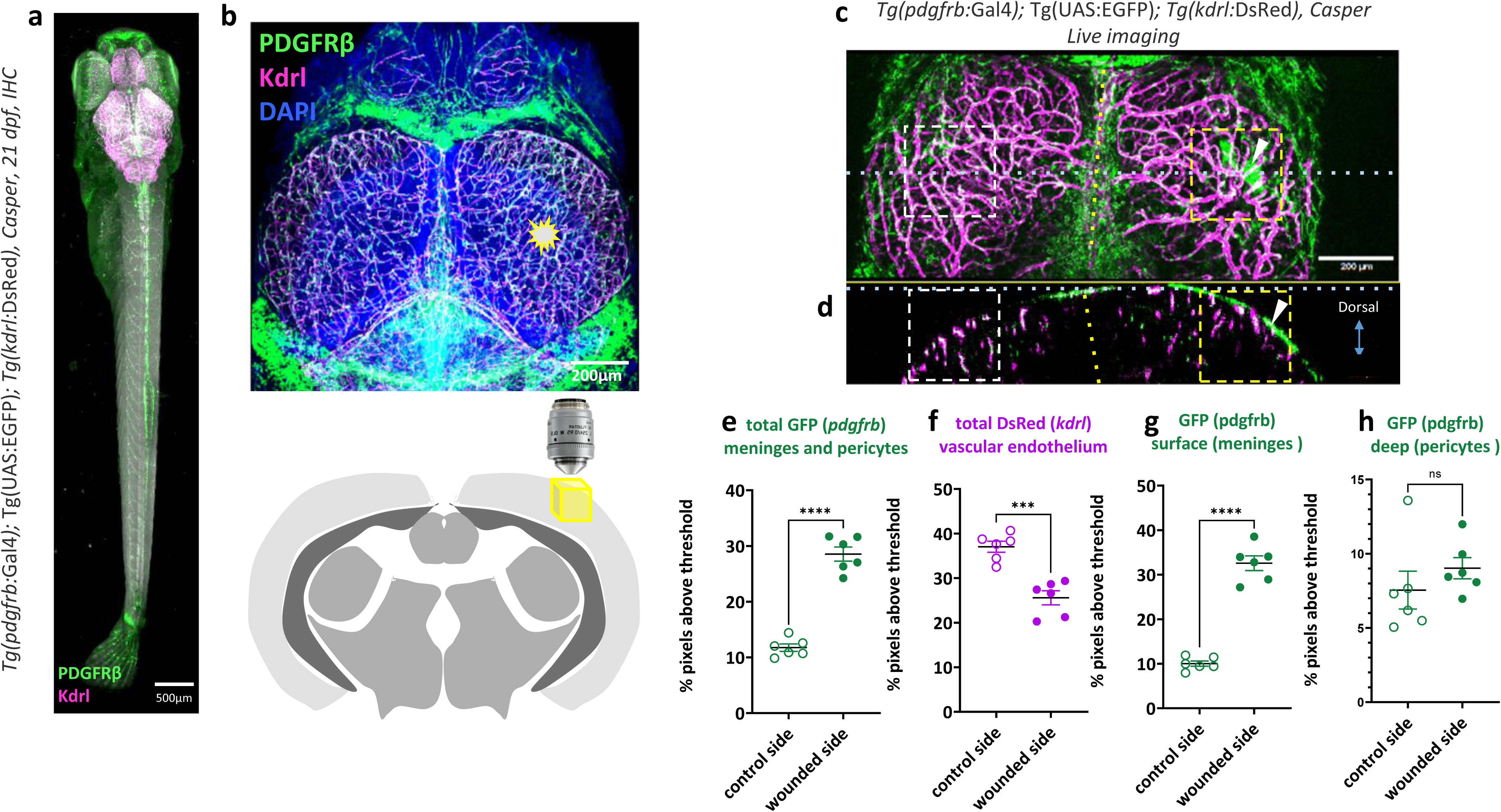
Recruitment of PDGFRβ^+^ meningeal cells to the wound following laser injury of the optic tectum. a, Immunostaining of a 21 days post-fertilization (dpf) *Tg(pdgfrb:*Gal4); Tg(UAS:EGFP)*; Tg(kdrl:*DsRed) zebrafish brain in a *casper* background. GFP⁺ *pdgfrb*-expressing cells (green) and *kdrl:DsRed*⁺ vasculature (red) are shown with DAPI nuclear counterstain (blue). b, Schematic of the two-photon laser ablation strategy targeting the optic tectum; the contralateral hemisphere serves as an internal control. The yellow box indicates the injury site. The schematic illustrates the objective lens orientation for the surface-localized laser injury. c, Intravital imaging at 24 hours post-injury (hpi) reveals the recruitment of GFP⁺ PDGFRβ^+^ meningeal cells (white arrowheads) to the wounded tectal surface, which are absent on the uninjured contralateral side. d, Orthogonal projection of the same sample confirms the localized accumulation of GFP⁺ meningeal cells at the injury site and their absence on the control side. e, Quantification of GFP expression (PDGFRβ^+^) shows a significant increase on the wounded side compared to control (****P < 0.0001). f, DsRed signal (kdrl:DsRed) marking vascular endothelium is significantly reduced at the wounded site (***P < 0.001). g, Quantification of GFP expression in surface-localized PDGFRβ^+^ meningeal cells reveal a robust increase in fluorescence intensity following injury (****P < 0.0001); h, In contrast, the deep pericyte-associated GFP signal remains unchanged (ns), indicating the selective recruitment of superficial PDGFRβ^+^ populations to the wound site.

### PDGFRβ signaling is required and sufficient for meningeal recruitment

To determine if PDGFR signaling is necessary for meningeal repair, we laser-wounded Tg(*pdgfrb:*Gal4; UAS:GFP*; kdrl:*DsRed) juveniles treated with the selective PDGFR inhibitor AG1295^28^ (Figure. 2a). At 24hpi, pharmacological inhibition of PDGFR completely abolished the injury-induced increase in GFP signal at the wound site (Figure. 2b). Spatial quantification revealed that this reduction was specifically restricted to the superficial meningeal layer, whereas PDGFRβ^+^ pericyte recolonization and vascular regrowth remained unaffected by AG1295 treatment (Figure. 2b). Our data indicate that PDGFR signaling is selectively required for the recruitment of superficial meningeal cells, but dispensable for pericyte-mediated vascular repair. While PDGFR inhibition significantly blocked the accumulation of arachnoid cells at the injury site, PDGFRβ^+^ pericytes successfully re-invaded the wound and associated with the regenerating vasculature (Figure. 2b, c). This striking divergence suggests that these two populations are functionally and mechanistically distinct. This observation is supported by recent multiome profiling, which reveals that arachnoid fibroblasts and pial/pericytic populations occupy separate transcriptional landscapes and utilize different molecular programs to respond to injury cues^35^. We next examined whether PDGFR activation is sufficient to drive meningeal accumulation. Following laser injury in *Tg(pdgfrb:Gal4; UAS:GFP)* fish, we observed the characteristic transient recruitment and subsequent decline of the meningeal signal at 4 dpi. Exogenous administration of recombinant human PDGF-BB, a validated agonist of zebrafish Pdgfrβ signaling via midline injection at 4 dpi triggered a robust and significant re-accumulation of GFP⁺ meningeal cells specifically at the wound site by 5 dpi (Figure. 2c,d). In contrast, control injections of BSA failed to elicit a response. Together, these loss- and gain-of-function experiments demonstrate that PDGFR signaling is both necessary and sufficient to orchestrate the targeted recruitment of meningeal cells to the site of CNS injury.

**Figure 2.**
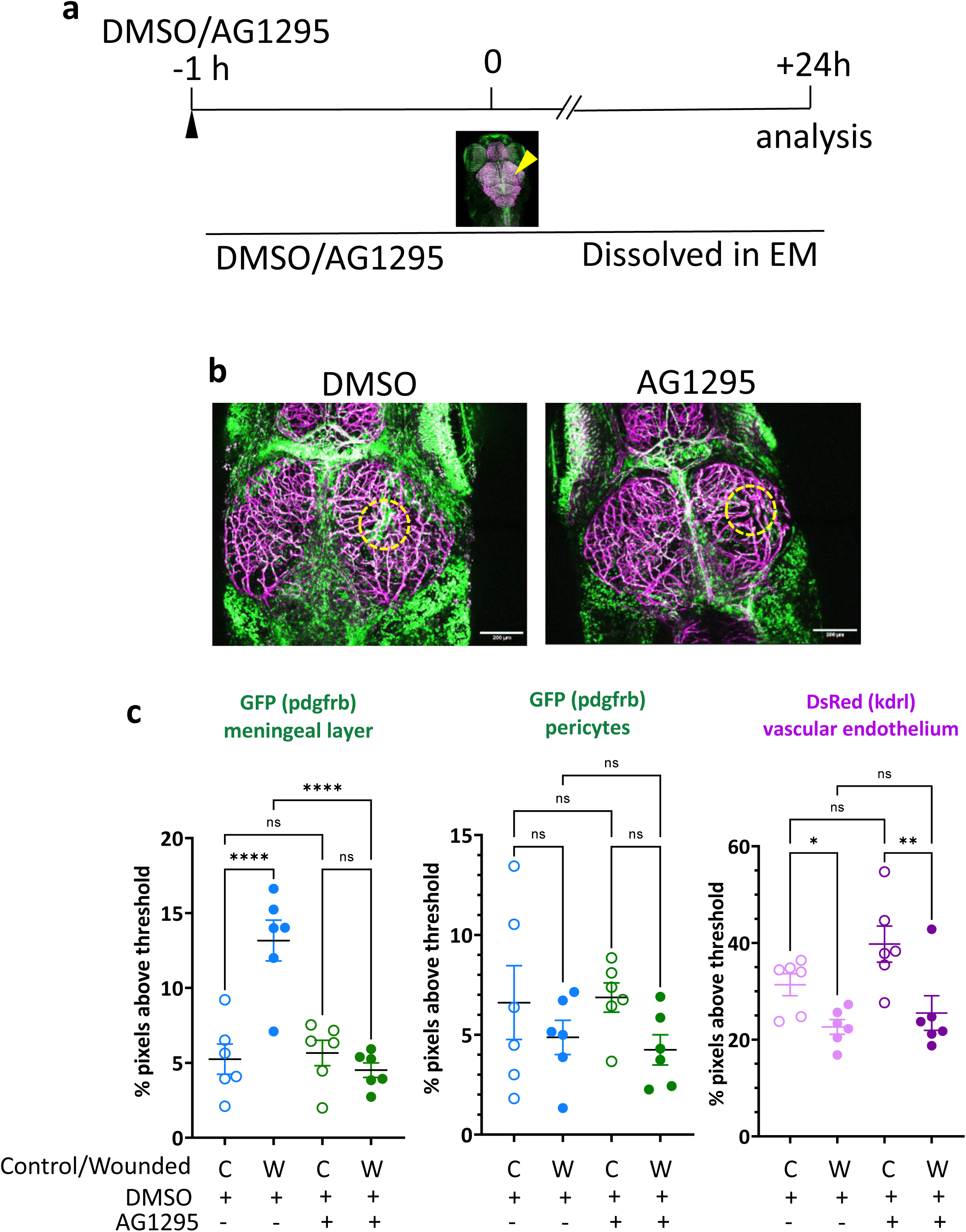
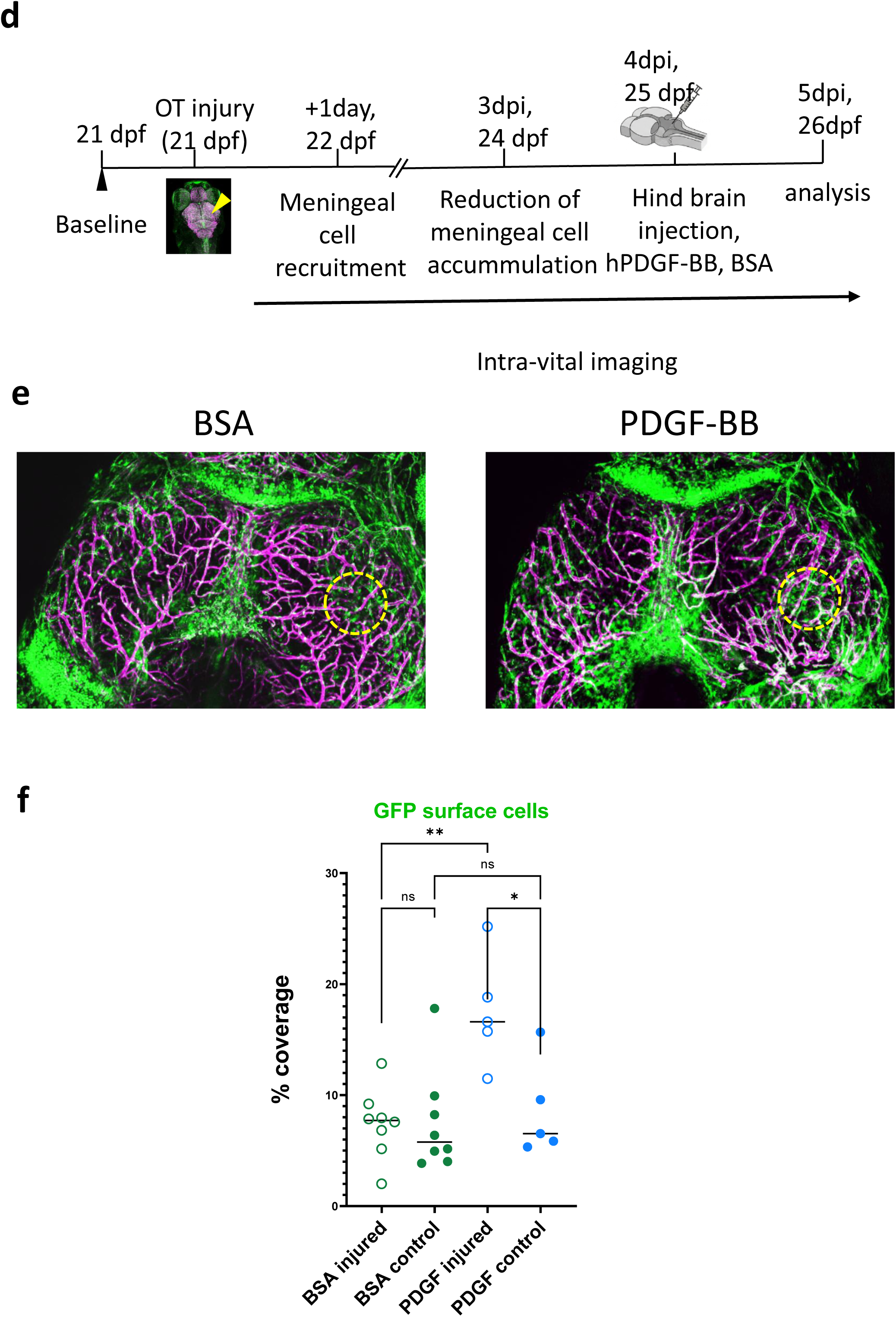
Pharmacological modulation of PDGFRβ+ meningeal cell response following injury. a, Schematic of the experimental timeline. Inhibitor/Vehicle was applied 1 hour prior to injury, followed by laser ablation at time 0 and analysis at 24 hours post-injury (hpi). b, Representative confocal images of Tg(*pdgfrb*:Gal4); Tg(UAS:GFP); Tg(kdrl:DsRed); casper zebrafish at 24 hpi, showing PDGFRβ^+^ cells (green) following treatment with vehicle (DMSO) or the PDGFR inhibitor AG1295 (30 µM). c, Left: Quantification of surface-localized GFP⁺ PDGFRβ^+^ cell coverage reveals a significant reduction in AG1295-treated larvae compared to vehicle-treated controls, indicating inhibition of injury-induced meningeal cell recruitment. Center: Quantification of deep pericyte-associated GFP signal shows no significant difference between treatment groups. Right: Kdrl⁺ endothelial cell coverage is further reduced on the injured side in AG1295-treated larvae compared to their contralateral control, with no significant difference observed between overall DMSO and AG1295 groups. d, Schematic representation of the experimental workflow from 21 to 26 days post-fertilization (dpf). Following baseline assessment at 21 dpf, a localized injury was performed on the optic tectum, initiating a period of meningeal cell recruitment at 1 dpi followed by a reduction in cell accumulation at 3 dpi. At 25 dpf 4 dpi, juveniles received a hindbrain microinjection of either PDGF-BB or BSA. Final analysis was performed at 26 dpf (5 dpi). Longitudinal intra-vital imaging was performed at 4 and 5 dpi to capture the response to pharmacological intervention. e, Representative confocal images showing PDGFRβ^+^cells (green) in juvenile with tectal injury. treated with PDGFR agonist hPDGF-BB (180 ng) or with vehicle (BSA). f, Quantification shows no significant difference in surface PDGFRβ^+^ cell coverage between injured and uninjured sides in BSA-treated controls. In contrast, PDGF-BB treatment significantly enhances pdgfrb⁺ cell recruitment beyond injury-induced levels.

To genetically confirm the requirement for PDGFR signaling with temporal precision, we employed the heat-shock inducible Tg*(hsp70l*:dn-pdgfrb-mCherry) line. In the absence of heat shock, transgenic larvae exhibited normal *pdgfrb*:GFP expression patterns, confirming the stability of the transgene under basal conditions (Figure S1, b). However, when heat shock was induced 6 hours prior to wounding to drive the expression of the dominant-negative receptor, the recruitment of GFP⁺ cells to the injury site was completely abolished (Figure S1, c). Notably, while pericyte-associated GFP expression remained detectable, the migration of the superficial meningeal population to the wound was specifically blocked. These results, mirroring our pharmacological data, provide definitive evidence that PDGFRβ signaling is the primary molecular driver of the meningeal regenerative response.

### PDGFRβ signalling modulates other aspects of surface optic tectum wound repair

We analyzed other aspects of wound repair after UV laser injury of the OT surface. Vessel regrowth was assessed using the vascular endothelium reporter Tg(*kdrl*:DsRed); as described previously, partial re-invasion of the injured area by not-yet luminized endothelial cells is observed at 24hpi; the vessel repair process is essentially complete by 7dpi^15^. As mentioned previously, quantification of DsRed+ cells coverage at 24hpi in presence of the PDGFRβ inhibitor did not reveal a significant difference with controls (Figure 2c).

Macrophages (including microglia) accumulate at injury sites and contribute to multiple aspects of repair. Here, we visualized these cells using the Tg(*mfap4*:mCherryF) transgenic line. At 24hpi, as expected, these cells congregated at the injury site (Figure 3a). Some mCherry-positive macrophages were seen in close proximity to GFP+ meningeal cells, but did not express GFP themselves. Treatment with the specific PDGFR inhibitor resulted in a significantly reduced macrophage accumulation but did not entirely abolish it (Figure 3b). This suggests that PDGFRβ^+^ meningeal cells act as a signaling hub. Recent single-cell characterization of the zebrafish meninges has identified specialized leptomeningeal cells with high secretory potential^36^. It is likely that the PDGFR-dependent recruitment of these cells is a prerequisite for the establishment of a pro-regenerative inflammatory environment.

**Figure 3.**
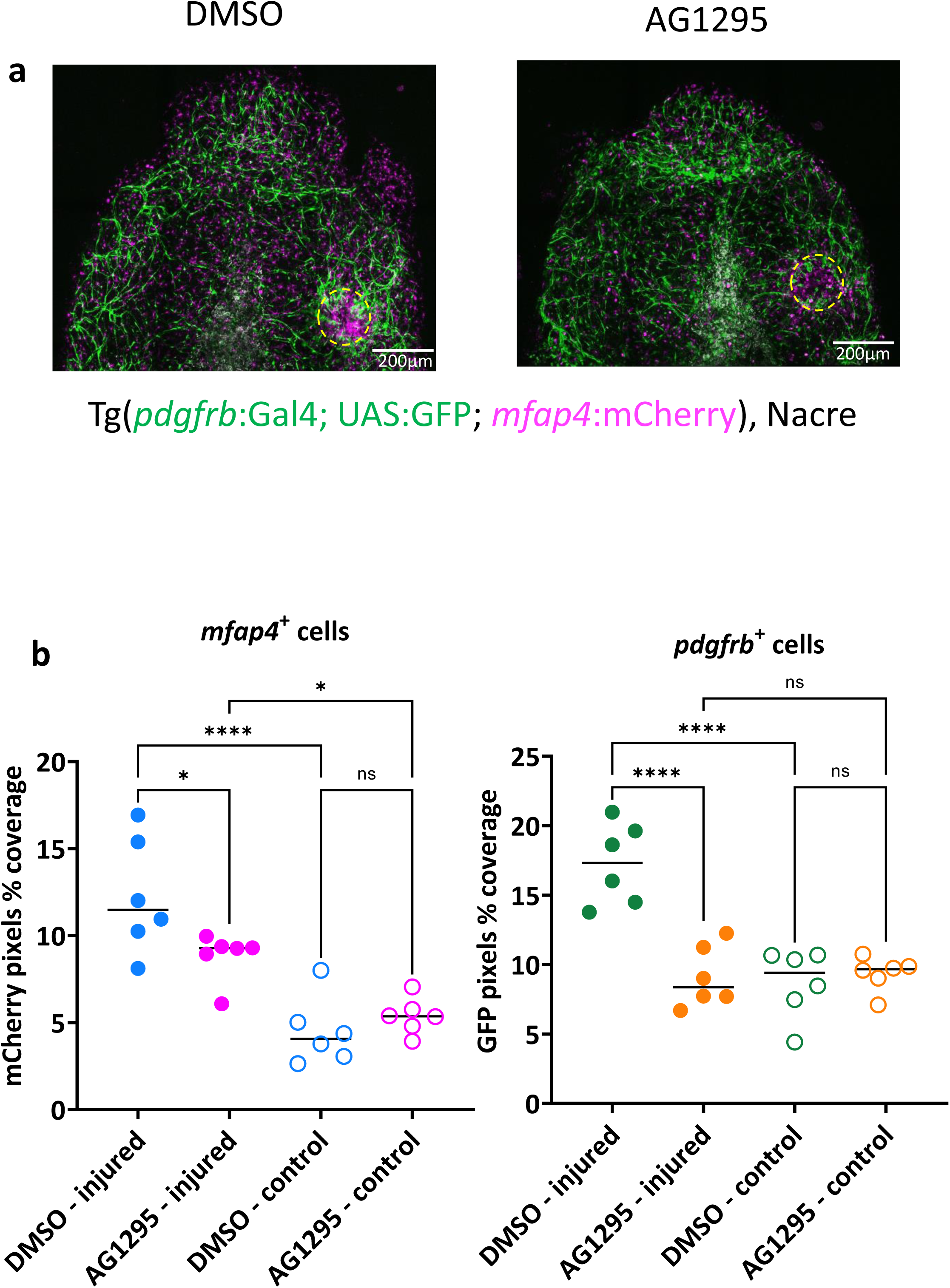
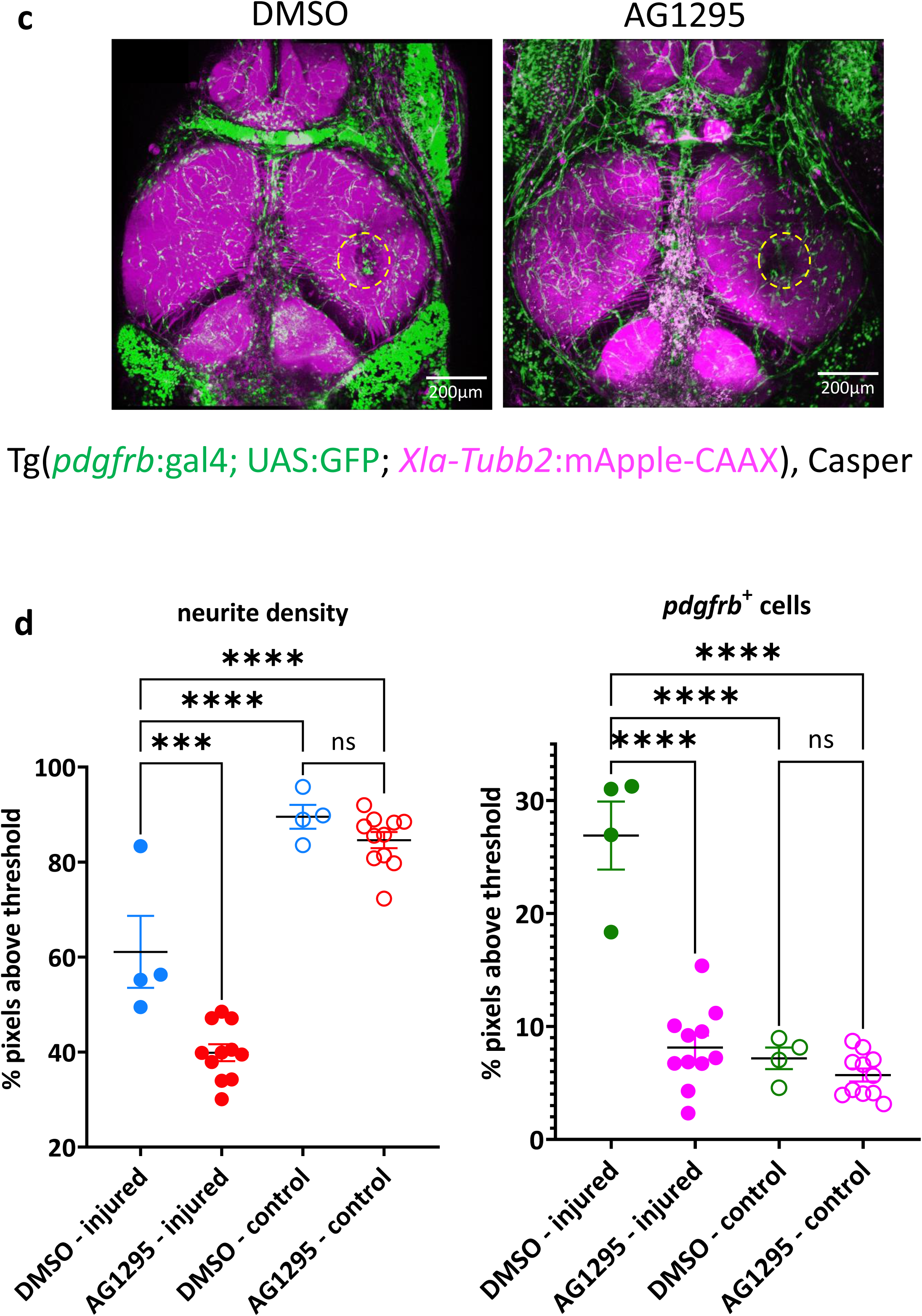
PDGFR inhibition impairs injury-induced accumulation of meningeal and microglial cells and reduces neurite density. a, Representative dorsal confocal images of Tg(*pdgfrb*:Gal4); Tg(UAS:GFP); Tg(*mfap4*:mCherry) zebrafish larvae treated with either vehicle (DMSO) or the PDGFR inhibitor AG1295 (30 µM), showing PDGFRβ⁺ meningeal cells (green) and mfap4⁺ macrophage-like cells (magenta) at the injury site (yellow dashed circle). DMSO-treated larvae exhibit substantial accumulation of mfap4⁺ cells at the wound, which is markedly reduced in AG1295-treated animals. b, Quantification of surface coverage by mfap4⁺ cells (left) and PDGFRβ^+^ cells (right). AG1295 treatment significantly reduces mfap4⁺ cell recruitment in the injured condition compared to DMSO (****p < 0.0001), with no effect in uninjured controls. A similar significant reduction in pdgfrb⁺ cell coverage is observed following AG1295 treatment (****p < 0.0001), with no change in controls (ns = not significant). c, Dorsal views of Tg(*pdgfrb*:Gal4); Tg(UAS:GFP); Tg(*Xla.Tubb2*:mApple-CAAX) larvae treated with DMSO or AG1295, showing neurite projections (magenta) and pdgfrb⁺ cells (green) at the injury site. d, Quantification of neurite density (left) and PDGFRβ^+^ cell surface coverage (right) in the injured hemisphere normalized to the contralateral side. AG1295 significantly reduces both neurite density (p < 0.001) and pdgfrb⁺ cell accumulation (****p < 0.0001) compared to DMSO-treated injured animals. No significant differences are observed between uninjured control groups.

Our biphoton-mediated wound procedure allows tight control of injury depth and damage to the brain parenchyma was essentially restricted to the upper layer of the optic tectum, a neuropile very rich in neurites but largely devoid of neuronal somata. To quantify the density of these neuronal processes, we used the Tg(*Xla.Tubb2*:mApple-CAAX) line, in which neurons express a membrane-addressed red fluorescent protein (Figure 3c). As expected, at 24 hpi, the fluorescence is reduced at the injury site in untreated animals. In presence of the PDGFRβ inhibitor, it was further diminished, implying that the regrowth of neurites into the damaged area is PDGFRβ-dependent (Figure 3d).

Finally, we checked for possible gross behavioral anomalies. Upon waking up from anesthesia after the laser injury, wounded larvae exhibited equilibrium defects, often laying on their side at rest, or swimming with occasional barrel rolls. However, at 4 hpi, all fish swam normally and maintained posture when immobile, indicating that this equilibrium defect is transient. At 24hpi, we measured spontaneous locomotion of control or injured fish incubated in presence or absence of the PDGFR inhibitor (Figure S2a). The distance travelled by the wounded fish was slightly reduced compared to controls, but no significant difference was found between DMSO and AG1295-treated fish. By 72hpi, no differences in velocity could be detected among groups (Figure S2b).

These findings establish that PDGFRβ^+^ meningeal cells function as a master regulatory hub during meningeal repair. Rather than acting in isolation, these cells orchestrate a multi-step regenerative cascade: they first mobilize to the lesion in a PDGFRβ dependent manner, subsequently triggering the recruitment of PDGFRβ^-^ immune populations and establishing the structural and molecular environment required for axonal regrowth. Our data thus position the meningeal response as the foundational event that dictates the success of both the inflammatory transition and neural regeneration.

### Spatiotemporal tracking of PDGFRβ^+^ cell origins

The morphology of the GFP⁺ clusters at the injury site is consistent with two distinct meningeal populations previously characterized^15^: the arachnoid cells resident within the forebrain–midbrain and midbrain–hindbrain junctions, and the dorsal midline population. While the latter is more challenging to resolve due to spectral interference from overlying autofluorescent pigment cells, both represent potential sources for the recruited population. While we previously hypothesized that the injury-responsive population derives from migrating arachnoid cells, the potential contribution of midline cells or the *de novo* upregulation of *pdgfrb* in GFP⁻ populations remained to be determined. To distinguish between these possibilities, we employed a Kaede-based spatiotemporal tracking assay using *Tg(pdgfrb:Gal4; UAS:Kaede)* larvae. Following laser injury, we selectively photoconverted (405 nm) either the midbrain–forebrain cleft arachnoid population or the midline population (Figure. S3).

The two-photon injury procedure induced localized photoconversion of pericytes immediately adjacent to the wound; however, this did not interfere with the analysis of the PDGFRβ⁺ meningeal response, as these cells are absent from the injury site at baseline. Despite partial reduction of the Kaede signal relative to the GFP reporter due to UAS-mediated silencing, the converted populations remained traceable over a 24-hour period. Selective targeting of arachnoid cells at the brain cleft resulted in the emergence of photoconverted (Kaede^red^) cells at the wound site in the majority of experimental replicates (n=4/6 animals; Figure. 4b). In contrast, photoactivation of the midline population yielded no converted cells at the wound (n=0/5 animals; Figure. 4c).

**Figure 4.**
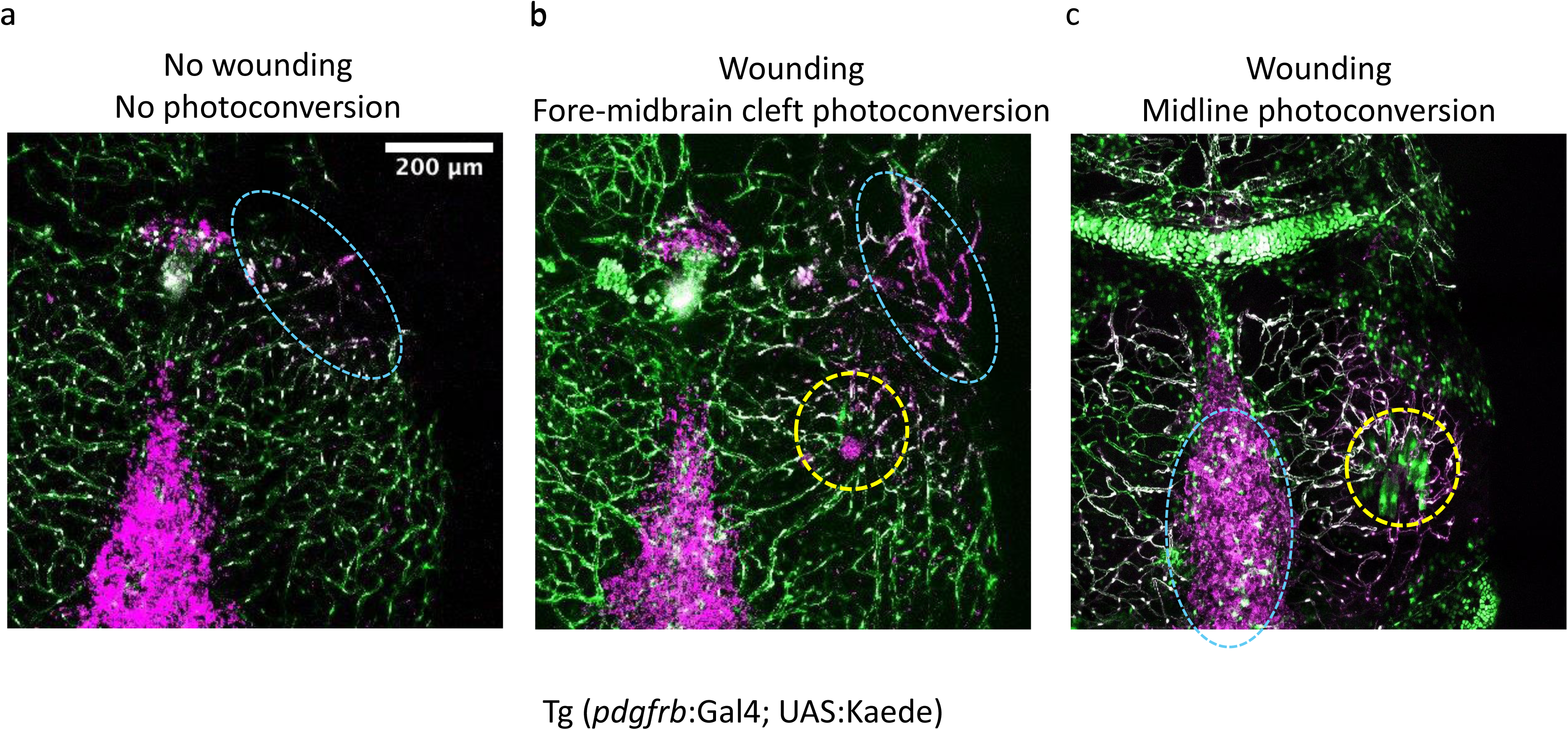
PDGFRβ^+^ meningeal cells migrating over the wound originate from the forebrain-midbrain cleft. a, representative Confocal image of a Tg(*pdgfrb*:Gal4); Tg(UAS:Kaede); nacre⁻/⁻ zebrafish larva showing photoconverted Kaede⁺ cells (magenta) in the forebrain-midbrain cleft (oval blue dashed) in the absence of wounding. No cell migration is observed toward the tectum. b, Following laser-induced tectal wounding (yellow dashed circle), photoconverted Kaede⁺ cells from the forebrain-midbrain cleft (blue dashed oval) are observed migrating toward the wound site. c, In contrast, when photoconversion is restricted to the midline (blue dashed oval) rather than the cleft, few or no photoconverted cells reach the wound area (yellow dashed circle),indicating the cleft as a primary source of wound-recruited PDGFRβ^+^ cells. Scale bar, 200 µm.

These lineage-tracing data suggest that the regenerative meningeal seal is primarily formed by the directed migration of resident arachnoid cells originating from inter-regional junctions. This finding identifies the brain fissures as significant anatomical reservoirs of repair-competent PDGFRβ^+^ cells that mobilize to restore the structural integrity of the meningeal barrier ^37,38^. While the lack of converted cells from the midline (0/5) and the arrival of cleft-derived cells (4/6) support this migratory model, the inherent limitations in photoconversion efficiency and sample size mean that localized de novo expression or minor contributions from other reservoirs cannot be definitively excluded. Nonetheless, the mobilization of these specific meningeal populations aligns with the emerging view of the meninges as a widespread niche for progenitor cells that react to central nervous system (CNS) injury^37,38^.

The proposed working model (Figure. 5) further suggests that injury-induced PDGFRβ signaling activates a FoxJ1-associated ciliary program, a mechanism previously identified through our single-cell RNA sequencing analysis of injured zebrafish^15^. This pathway likely enhances ciliary function and migratory competence, facilitating the rapid mobilization of these reservoir cells to the site of injury to restore the meningeal barrier.

**Figure 5.**
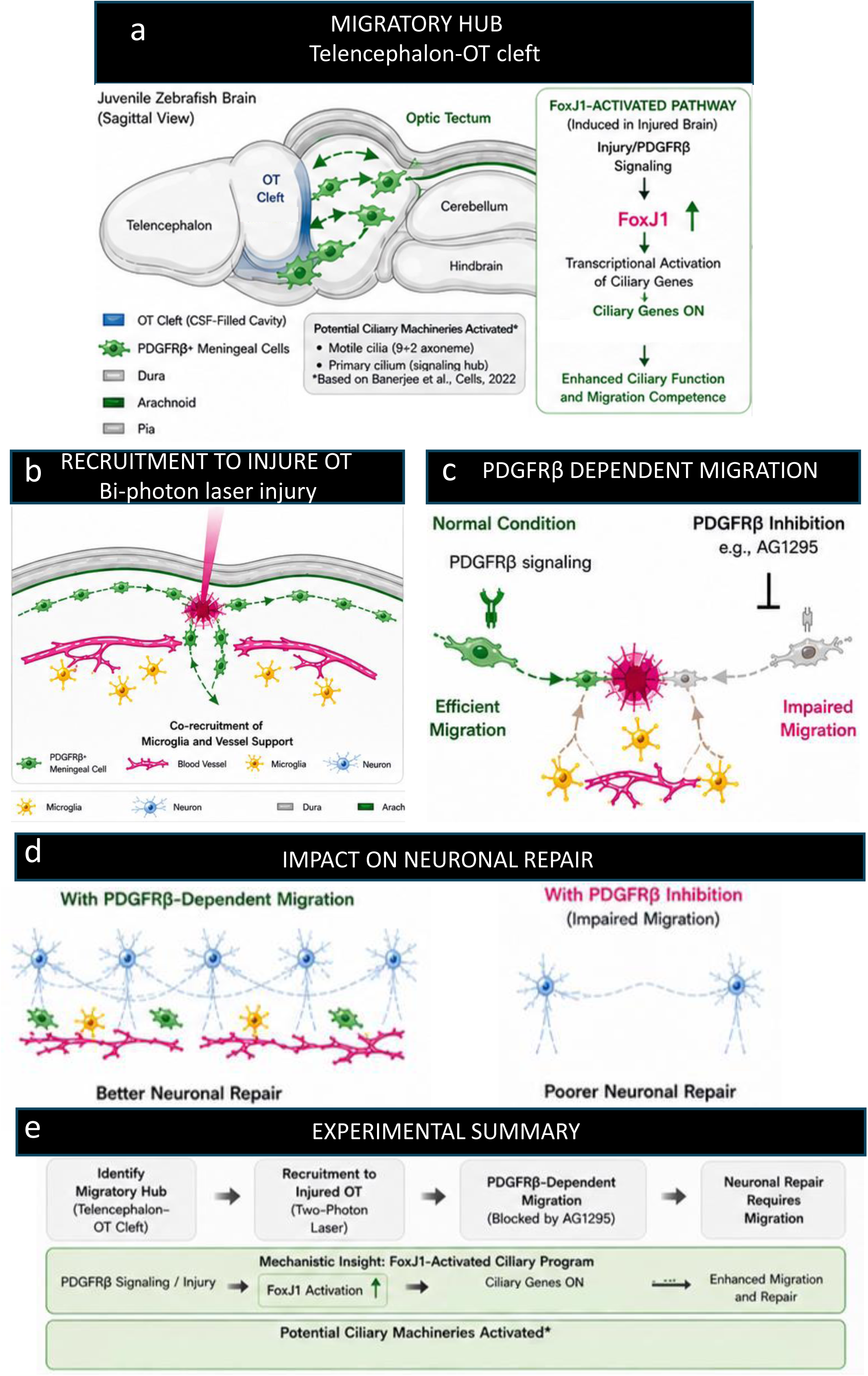
PDGFRβ signaling drives meningeal cell migration and promotes meninges repair. Schematic summary illustrating the proposed mechanism of PDGFRβ-dependent meningeal cell migration following optic tectum (OT) injury in juvenile zebrafish. (a) A migratory hub of PDGFRβ^+^meningeal cells is localized near the telencephalic midline/OT cleft region in the sagittal brain. (b) Following two-photon laser injury to the optic tectum, meningeal cells are recruited toward the lesion site along the meningeal surface. (c) Under normal conditions, PDGFRβ signaling supports efficient migration toward the injured region, whereas pharmacological inhibition of PDGFRβ signaling (e.g., AG1295 treatment) impairs migratory responses. (d) Efficient meningeal migration is associated with improved neuronal repair outcomes, while inhibition of migration correlates with reduced repair efficiency. (e) The experimental summary highlights that neuronal repair is fundamentally dependent on the recruitment of meningeal cells from the Telencephalon-OT Cleft migratory hub. Following two-photon laser injury, PDGFRβ signaling triggers a mechanistic program characterized by FoxJ1 activation and the transcriptional upregulation of ciliary genes. This FoxJ1-activated ciliary program enhances ciliary function potentially utilizing motile and primary ciliary machineries to provide the migration competence necessary for efficient cellular movement and successful neuronal restoration.

## Discussion

Following the unexpected finding of flat PDGFRβ*⁺* cells appearing over a wound of the optic tectum of juvenile zebrafish in our previous study, we explored the mechanisms of meningeal repair in this optically tractable model. Our data support a model (Figure. 5) in which meningeal injury triggers the mobilization of PDGFRβ⁺ arachnoid-like cells from specialized reservoirs in the forebrain-midbrain junction clefts. These cells migrate to the lesion site to reconstruct the brain’s protective barrier, forming a “seal” that acts as a master regulatory hub. The re-establishment of this meningeal layer is a prerequisite for the subsequent recruitment of immune cells and the promotion of robust axonal regrowth. Our results establish that PDGFRβ signaling is not merely a marker of arachnoid cells but a functional requirement for their mobilization. This aligns with recent findings that emphasize PDGFRβ as a master regulator of the ‘regenerative niche’, coordinating the transition from an inflammatory to a pro-regenerative state^35^. By acting as a molecular bridge, PDGFR activity ensures that the structural repair of the meninges is synchronized with cellular influx^39^. Crucially, this repair mechanism likely intersects with established pathways of vertebrate CNS regeneration. Our previous transcriptomic analysis revealed that *foxj1* levels are induced in these cells following injury^15^. This is highly significant given that in other zebrafish CNS models, Foxj1a^+^ cells expand in response to injury through a Sonic Hedgehog (Shh)-dependent mechanism to restore tissue architecture^40^. We propose that a similar Shh/Foxj1 axis may operate downstream of, or in parallel with, PDGFRβ signaling to grant these arachnoid cells the migration competence and “ciliary machinery” required to seal the brain wound. These cells are morphologically very distinct from pericytes, of which PDGFRβ is also a well-known marker^41, 42^. Pericytes are abundant on the borders of blood vessels in the parenchyma but largely absent from meninges; by performing depth-selective analysis of our 3D-images of live zebrafish, we could separately quantify the abundance of the meningeal and pericytes populations, despite sharing the same transgenic marker. Thus, we established that the robust increase of GFP signal over the wounded area in Tg(*pdgfrb*:Gal4; UAS:GFP) zebrafish was due to a strong increase of GFP+ meningeal cells rather than pericytes, although the latter also re-invade the wounded area. To address the role of PDGFRβ in this meningeal repair mechanism, we used a selective pharmacological inhibitor, AG1295^7,28^. Addition of this inhibitor totally prevented the appearance of the GFP+ meningeal cells, but had no measurable impact on the recolonization of the wounded parenchyma by pericytes or endothelial cells. These meningeal PDGFRβ⁺ cells were also found to re-increase in numbers in response to injection of PDGFBB^7,28^, but only over the previously wounded area; this is consistent with their strong dependence of PDGFRβ, and implies that other signals from the wound control their influx. We attempted to confirm these results using a genetic approach based on a heat-shock inducible transgene encoding a dominant-negative *pdgfrb* fused to mCherry^2^. Our results with a dominant-negative (DN) *pdgfrb* transgene further support this; the pre-existing absence of arachnoid cells in carrier fish prevented any subsequent wound accumulation. This differential sensitivity suggests that arachnoid cells require a specific PDGFRβ signaling threshold for maintenance that is higher than that required by pericytes. Using the PDGFRβ inhibitor, we could also measure a small but significant reduction of the accumulation of macrophages/microglia at the wounded site, and a reduction of neurite regrowth. Since macrophages and neurons express very little if any *pdgfrb*, we hypothesize that this is an indirect effect of the perturbed meningeal repair that we observe with PDGFRβ inhibition. Alternatively, pericytes re-invading the wound may also secrete less chemoattractants and neurite-growth promoting factors if they receive no PDGFRβ signal, while retaining their recolonization properties. Whatever the reason, it is interesting to observe that in the well-characterized larval spinal cord resection model, axonal regrowth was similarly dependent on PDGFRβ, but macrophage accumulation was not^2^. Since a key difference between these two CNS regeneration models is the presence of PDGFRβ⁺ cells, we hypothesize that PDGFRβ⁺ arachnoid cells may produce chemokines to attract more macrophages to the wound. Despite these various measurable effects on wound repair, we could not measure a significant difference in the locomotion (or other behavioral parameters, such as thigmotaxis or shoaling, not shown) of juvenile fish in presence or absence of the inhibitor, despite a measurable reduction caused by the wound itself. This may not be surprising considering the role of the optic tectum in processing visual signals, with no direct control of spontaneous locomotion. Another important finding of our work is the demonstration, using photoconversion, that PDGFRβ⁺arachnoid cells located in the clefts bordering the midbrain migrate towards the wound and thus participate in meninges repair, confirming the hypothesis from our previous work^15^. We presume that all the GFP+ cells that we detect over the wound at 24hpi derive from this population, although we cannot rule out de novo expression of *pdgfrb* by some previously negative meningeal cells or precursors thereof. It will be an interesting line of research in the future to identify the signals from the wound that mobilize these cells besides PDGFRβ ligands, and if similar mechanisms operate in mammals.

While our study identifies PDGFRβ⁺ arachnoid cells as the primary drivers of meningeal repair, the cellular landscape of the zebrafish meninges remains more complex and less characterized than its mammalian counterparts. Specifically, a unique population of single, non-vessel-forming cells related to the lymphatic endothelium has been identified within the meningeal layers^12^. These cells express classical lymphatic endothelial markers but are distinguished by their lack of *pdgfrb* expression under homeostatic conditions. Furthermore, our transcriptomic analysis does not suggest that these cells acquire *pdgfrb* expression following injury^15^, reinforcing the idea that they represent a lineage distinct from the migratory arachnoid cells characterized here. Nevertheless, investigating the behavior of these lymphatic-related cells during the wounding response remains a priority for future research. Understanding whether they actively interact with or provide secondary cues to the migrating PDGFRβ⁺ arachnoid population could yield deeper insights into the integrated cellular crosstalk required to successfully reconstruct the brain’s protective barrier.

In conclusion, our findings reveal that PDGFRβ, already known for its contribution in a variety of tissue repair models via its action on pericytes and fibroblasts, also plays a key role in meningeal repair by promoting the migration of distant arachnoid cells over the wound. Our findings underscore that PDGFRβ signaling, acting through a downstream Foxj1-activated ciliary program, is a primary driver of meningeal repair. This axis orchestrates the synchronization of structural barrier reconstruction with the cellular crosstalk essential for CNS regeneration. These insights offer new therapeutic avenues for treating brain trauma and provide a mechanistic framework for understanding meningioma invasiveness, wherein PDGFRβ and Foxj1 signaling may be co-opted to drive cellular motility.

## Data availability

The datasets generated and analyzed during the current study are available from the corresponding authors upon reasonable request. A complete data package, including raw imaging files and processed quantification sheets, is currently being curated and will be deposited in a public repository (Zenodo) upon submission to a peer-reviewed journal.

## Acknowledgements

This manuscript is fondly dedicated to the memory of Jean-Stéphane Joly, who initiated this project. We are indebted to Matthieu Simion, Fabrice Licata and Pithursan Karunathasan who contributed to set up the imaging pipeline during their stay at TEFOR Paris-Saclay. Luc Jouneau (INRAE, Jouy-en-Josas) provided valuable feedback and training on our datasets. Daniel Wehner (Max Planck Institute, Erlingen, Germany) provided transgenic zebrafish lines and precious advice. Jean-Paul Concordet (TACGene, MNHN, Paris) provided the SpCas9 protein. The work presented here is the continuation of a project initiated by funds from the CELPHEDIA French National Infrastructure for research and development.

## Author contributions

P.B. and J.-P.L. conceived the study, designed the experimental framework. P.B. developed the methodologies and performed the majority of the experimental assays, with technical assistance from M.P. regarding data collection. M.D. and A.B. optimized and executed the PDGF-BB intracranial injections. A.B. performed the CRISPR/Cas9 mutagenesis, while D.C. conducted the whole-mount imaging. A.J. optimized the 3D segmentation and produced the 3D movies. C.F. and H.W. assisted P.B. with the locomotion assays. Data analysis was conducted by P.B., M.D., and J.-P.L. The manuscript was drafted by P.B., J.-P.L. M.D. and A.J. contributed to the final review and editing of the text. J.-P.L. secured funding and supervised the project. P.B. and J.-P.L. serve as corresponding authors and contributed equally to the coordination of this work.

---

**Supplementary Movie 1. 3D spatial organization of recruited PDGFRβ^+^ meningeal cells.**

3D reconstruction of a 21 dpf *Tg(pdgfrb:*Gal4); Tg(UAS:EGFP); Tg(kdrl:DsRed) brain in a *casper* background following injury. PDGFRβ^+^ cells (green) and *kdrl*⁺ endothelial cells (magenta) are shown. Recruited *pdgfrb*⁺ cells localized to the injury site in the left midbrain are segmented and pseudo-colored in yellow; these cells exhibit an arachnoid-like morphology and are absent from uninjured regions. Spatially distinct, non-pericytic PDGFRβ^+^ populations located at the forebrain–midbrain and midbrain–hindbrain boundaries, as well as the midline, are segmented and pseudo-colored in blue.

**Figure S1. Inhibition of PDGFRβ signaling impairs meningeal cell accumulation following injury.**

a, Schematic of the experimental timeline. Heat shock (HS) was applied 6 hours prior to injury (–6 h), followed by laser ablation at time 0 and analysis at 24 hours post-injury (hpi).

b, Representative dorsal confocal image of a Tg(*pdgfrb*:Gal4); Tg(UAS:EGFP); Tg(*hsp70l:dn-pdgfrb*-mCherry) zebrafish larva without heat-shock treatment, showing robust accumulation of PDGFRβ⁺ meningeal cells (green) at the injury site (yellow dashed circle).

c, Representative dorsal confocal image of a Tg(*pdgfrb*:Gal4); Tg(UAS:EGFP); Tg(*hsp70l:dn-pdgfrb*-mCherry) larva following heat-shock treatment, as depicted in the experimental schema (a), showing co-expression of EGFP⁺ meningeal cells (green) and dominant-negative PDGFRβ -mCherry (magenta). Impaired PDGFRβ signaling markedly reduces the recruitment of pdgfrb⁺ cells to the wound site (yellow dashed circle).

**Figure S2. PDGFR inhibition does not impair recovery of locomotor activity following surface brain injury.**

a, Representative swim trajectories of zebrafish larvae at 24 hours post-injury (hpi) and in uninjured controls, following treatment with either vehicle (DMSO) or the PDGFR inhibitor AG1295 (30 µM). Injury leads to reduced locomotor activity at 24 hpi in both AG1295- and DMSO-treated larvae. No overt differences are observed between treatments under uninjured conditions.

b, Quantification of distance traveled (cm/min) at 24 hpi (left) and 72 hpi (right). At 24 hpi, AG1295-treated injured larvae exhibit significantly reduced locomotor activity compared to uninjured AG1295-treated controls (**p < 0.01, ****p < 0.0001). No significant differences are detected between AG1295- and DMSO-treated groups within the same injury condition. By 72 hpi, locomotor activity is restored across all groups, indicating full recovery. Data are presented as mean ± SEM. Statistical analysis was performed using one-way ANOVA followed by post hoc testing. ns, not significant.

**Figure S3. Successive laser wounding of the tectum and photoconversion of arachnoid cells in the forebrain-midbrain cleft.**

a, Schematic timeline of the photoconversion assay in juvenile zebrafish. Subsequent live imaging and cell fate analysis were conducted at 22 dpf (1 day post-injury, dpi; T=+24h) to track the migration and lineage of photoconverted cells during the inflammatory response.

b, Confocal image showing unconverted Kaede-expressing perivascular or meningeal fibroblast-like cells (green) in the tectum of a Tg(*pdgfrb*:GAL4; UAS:Kaede); nacre⁻/⁻ zebrafish larva prior to wounding. Magenta signal along the dorsal midline corresponds to autofluorescence from xanthophore pigments.

c, Same region imaged after laser-induced tectal wounding (yellow dashed circle); Kaede remains unconverted, and xanthophore autofluorescence is retained.

d, Following photoconversion in the forebrain-midbrain cleft (blue dashed oval), Kaede-expressing cells exhibit red fluorescence. The wound site remains marked (yellow dashed circle).

