## Supplementary figures and images for "Pdgfrβ signaling orchestrates meningeal repair via the mobilization of arachnoid cells"

### Supplemental Figure 1

Figure S1

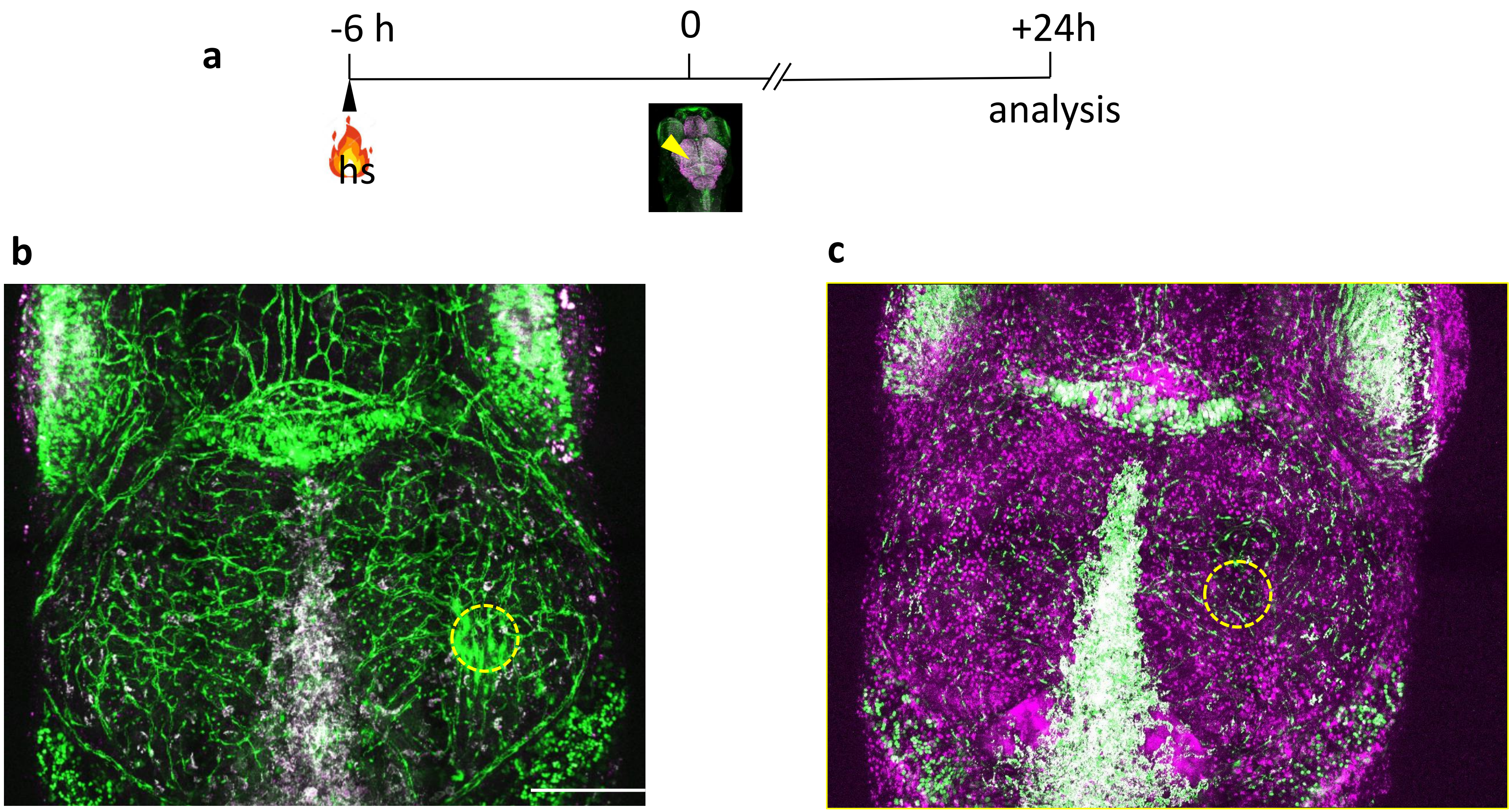


*pdgfrb*:Gal4; UAS:EGFP

*pdgfrb*:Gal4; UAS:EGFP;
*hsp70ldnpdgfrβ*-mcherry

### Supplemental Figure 2

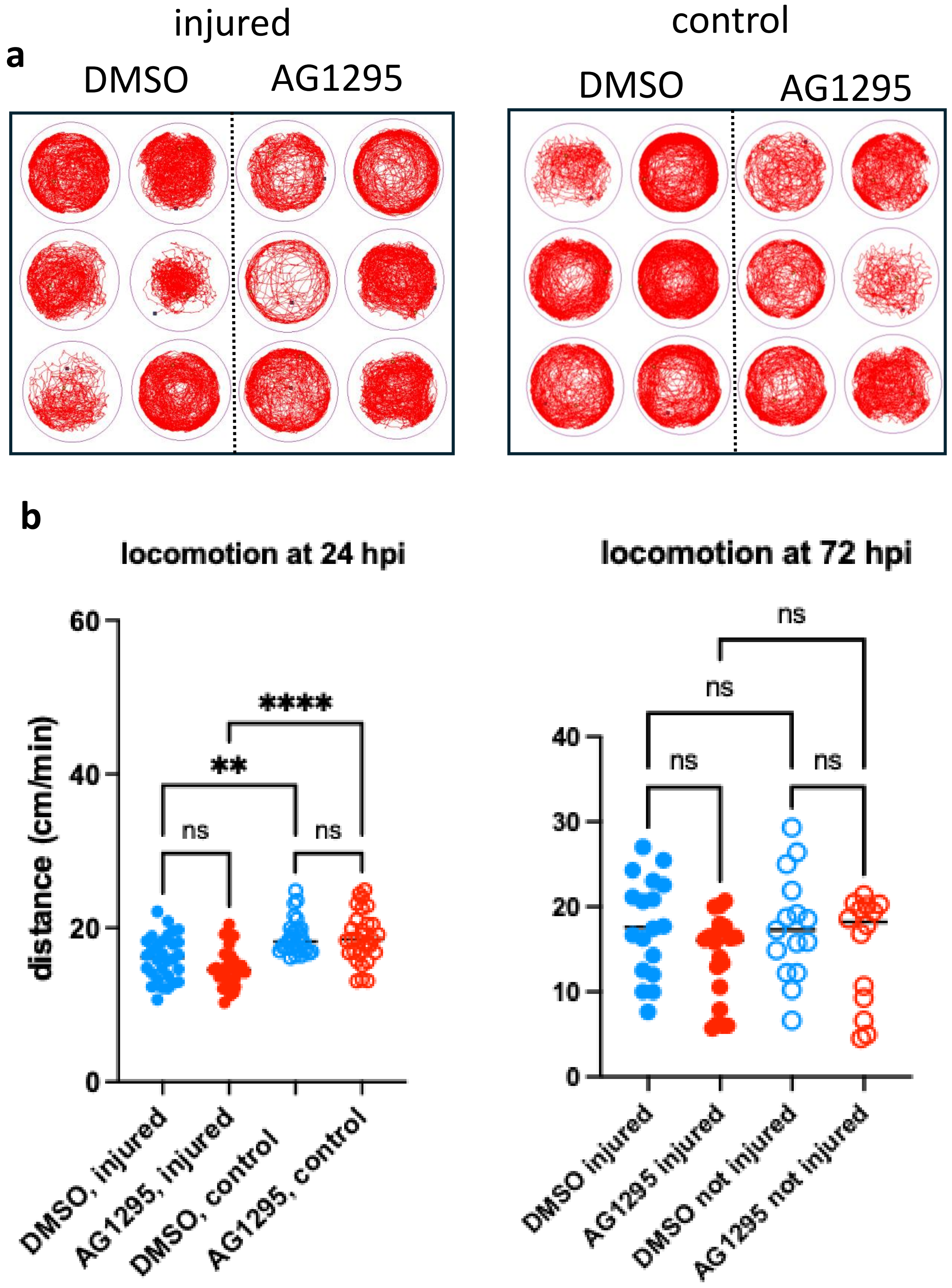

### Supplemental Figure 3

Figure S3

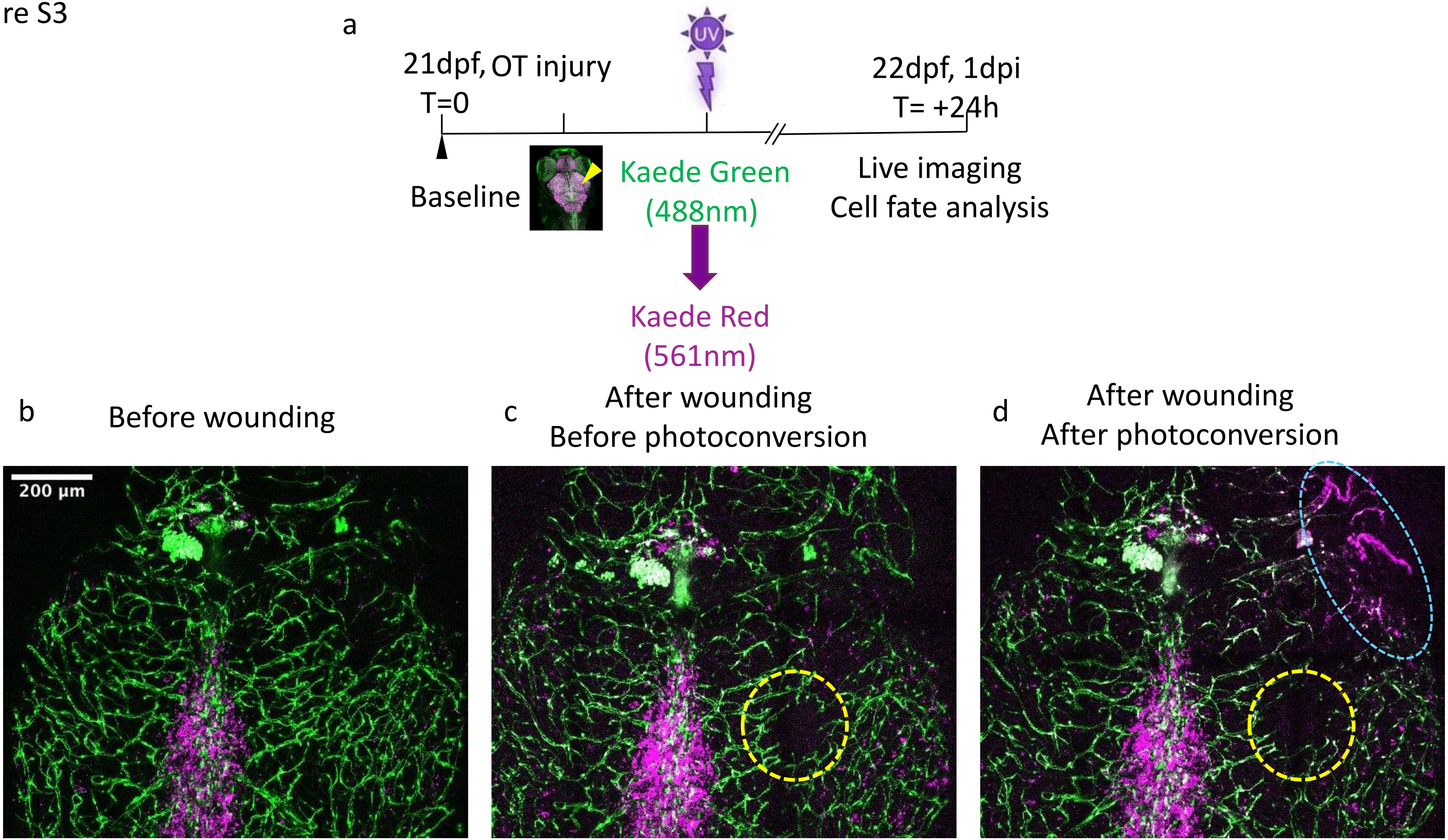


Tg (*pdgfrb*:Gal4; UAS:Kaede), Nacre
